# The molecular clockwork of the suprachiasmatic nucleus is sufficient to co-ordinate phasing and stabilisation of sleep-wake cycles and enhance memory deficits in a clockless mouse

**DOI:** 10.1101/2021.02.04.429717

**Authors:** Elizabeth S. Maywood, Johanna E. Chesham, Raphaelle Winsky-Sommerer, Michael H. Hastings

## Abstract

The timing and quality of sleep-wake cycles are regulated by interacting circadian and homeostatic mechanisms. Although the suprachiasmatic nucleus (SCN) is the principal circadian clock, local clocks are active across the brain and the respective sleep-regulatory roles of SCN and extra-SCN clocks are unclear. To determine the specific contribution(s) of the SCN, we used virally mediated genetic complementation, expressing Cryptochrome1 (Cry1) to restore circadian molecular competence to the SCN of globally clockless *Cry1/Cry2*-null mice. Under free-running conditions, the rest/activity behaviour of *Cry1/Cry2*-null controls which received EGFP (SCN^Con^) was arrhythmic, whereas Cry1-complemented mice (SCN^Cry1^) had circadian behaviour comparable to that of Cry1,2-competent wild-types (WT). In SCN^Con^ mice, sleep-wakefulness, assessed by electroencephalography/electromyography, also lacked circadian organisation. In SCN^Cry1^ mice, however, it was comparable to WT, with consolidated vigilance states (wake, REM and NREM sleep) and rhythms in NREMS delta power and expression of REMS within total sleep. Wakefulness in SCN^Con^ mice was more fragmented than in WT, with more wake-NREMS-wake transitions. This disruption was corrected in SCN^Cry1^ mice. Following sleep deprivation, all mice showed an initial homeostatic increase in NREMS delta power. The SCN^Con^ mice, however, had reduced, non-consolidated NREMS during the inactive phase of the recovery period. In contrast, the dynamics of homeostatic responses in the SCN^Cry1^ mice were equivalent to WT. Finally, SCN^Con^ mice exhibited poor sleep-dependent memory but this was corrected in SCN^Cry1^mice. Therefore, the SCN clock is sufficient for circadian control of sleep-wake, facilitating initiation and maintenance of wake, promoting sleep consolidation, homeostatic dynamics, and sleep-dependent memory.

**Significance statement:** The circadian timing system regulates sleep-wake cycles. The hypothalamic suprachiasmatic nucleus (SCN) is the principal circadian clock, but local clocks are also active across the brain and the respective roles of SCN and local clocks in regulating sleep are unclear. To determine, explicitly, the contribution of the SCN, we used virally mediated genetic complementation to restore SCN molecular circadian functions in otherwise genetically clockless mice. This initiated circadian activity-rest cycles, accompanied by circadian sleep-wake cycles, circadian patterning to the intensity of NREM sleep and circadian control of REM sleep as a proportion of total sleep. Consolidation of sleep-wake established normal dynamics of sleep homeostasis and enhanced sleep-dependent memory. Thus, the SCN is the principal and sufficient circadian regulator of sleep-wake.

## Introduction

The timing and quality of sleep are determined by a circadian process that ensures sleep occurs appropriately within the light-dark (LD) cycle and a homeostatic process that tracks sleep need during wakefulness (Borbely and Achermann, 1999; Borbely et al., 2016). Whereas the identity of the homeostat remains unknown, the hypothalamic suprachiasmatic nucleus (SCN), is conventionally thought to mediate circadian control (Saper et al., 2005). At the molecular level, the SCN clock consists of transcriptional/post-translational feedback loops (TTFL) in which *Period* (*Per*) and *Cryptochrome* (*Cry*) genes are trans-activated by CLOCK and BMAL1 heterodimers. Following their accumulation over circadian day, the encoded Per and Cry proteins inhibit trans-activation, closing the loop, whilst their subsequent degradation over circadian night allows the cycle to recommence. This TTFL is entrained to solar time by melanopsin-containing retinal ganglion cells that innervate the SCN (Tsai et al., 2009). In constant darkness (DD), however, it runs to its intrinsic approximately 24h period. Importantly, the TTFL is active in all tissues, including brain regions that regulate sleep/wake and memory, which are synchronised by the SCN (Hastings et al., 2018). The question arises, therefore, as to whether circadian control of sleep is mediated uniquely by the SCN, or do local brain clocks also contribute? Beyond that, what influences does the circadian system (SCN and/or local clocks) have over sleep-wake cycles: does it only affect timing, or does it modify their temporal architecture or homeostatic responses (Gillette, 2004)? Intriguingly, *Per2* responds to sleep loss and time of day, and so is well positioned to bridge circadian and homeostatic controls (Curie et al., 2013).

Loss-of-function ablation has demonstrated the necessity of the SCN for circadian timing of sleep under DD, and its redundancy in global sleep homeostasis (Tobler et al., 1983; Mistlberger, 2005), although increased NREM sleep (NREMS) in SCN-ablated mice suggests a broader role in sleep regulation (Easton et al., 2004). Loss-of-genetic function approaches have also been used to interrogate circadian sleep control. *Per1/Per2*-null mice have a defective TTFL and arrhythmic sleep-wake patterns under DD, but not LD (Shiromani et al., 2004). Moreover, sleep deprivation induces expression of *Per1* and *Per2* in the cerebral cortex (Franken et al., 2007), but *Per1/Per2*-null mice show normal homeostatic regulation of the daily amounts of waking, NREMS, or REM sleep (REMS) (Shiromani et al., 2004). Similarly, *Cry1/Cry2*-null mice also lose TTFL function and so have no circadian pattern to sleep-wake on DD but do show higher levels of NREMS and delta power (Wisor et al., 2002). Further, when sleep-deprived under a LD schedule, *Cry1/Cry2*-null mice show less NREMS recovery than do WT mice. This may indicate a requirement for Cry proteins in sleep homeostasis, and/or it may be a consequence of the prior increased NREMS and delta power in *Cry*-null mice. Finally, CLOCK mutant mice show altered sleep homeostasis (Naylor et al., 2000).

Untangling the anatomical (SCN, extra-SCN) and genetic (*Per, Cry, Clock*) contributions to sleep regulation has therefore been challenging. Interpretation of results from clock gene mutants is constrained because global mutations compromise both the SCN and local clocks. Furthermore, as transcription factors, their encoded proteins may have non-circadian roles. Equally, SCN ablations disrupt neural circuitry and may compromise non-circadian processes (Mistlberger, 2005). As an alternative, therefore, we employed a gain-of-function approach. Behavioural arrhythmia in clock mutant mice can be rescued by SCN grafting (Sujino et al., 2003) or by virally mediated genetic complementation of the SCN (Fuller et al., 2008; Maywood et al., 2018). But what of sleep regulation? Is a molecularly competent SCN clock sufficient to co-ordinate phasing and stabilisation of sleep-wake cycles in an otherwise clockless and arrhythmic mouse? We used adeno-associated viral (AAV) vectors to express control EGFP or a Cry1::EGFP fusion in the SCN of *Cry1/Cry2*-null mice. Leaving local clocks dysfunctional, would this be sufficient to co-ordinate phasing and stabilisation of sleep-wake cycles and, if so, would it enhance sleep-dependent memory in an otherwise clockless mouse?

## Materials and Methods

### Animals and Housing

All experiments were conducted in accordance with the UK Animals (Scientific Procedures) Act of 1986, with local ethical approval (MRC LMB, AWERB). We used 4-6 months-old male WT mice and *Cry1, Cry2* double knock-out mice (CDKO) (van der Horst et al., 1999), all on a C57/BL/6J genetic background. There were no significant differences in body weights at the time of surgery (WT =30.5 ±1.6g; CDKO =28.5 ±1.1g, n =5, 18 respectively). Mice were housed individually and their activity patterns were monitored continuously using running-wheels. Food and water were provided *ad libitum*. Mice were entrained to a 12h:12h light:dim red light cycle (LD) for at least 10 days before transfer to a schedule of continuous dim red light (DD) for 14 days for assessment of (ar)rhythmicity (DD1). Following surgery (see below), mice were maintained on a 12L:12D photoschedule for recovery before transfer to a second period of DD (DD2). In all of our studies, Zeitgeber time (ZT) 0 denotes the time of lights-on and ZT12 lights-off under LD, whereas circadian time (CT) 0 denotes the start of subjective day and CT12 denotes the start of subjective night in DD, as evidenced by activity onset.

### Stereotaxic injection of AAV vector and implantation of EEG/EMG transmitters

Mice were anaesthetised using isoflurane (induction 2-4%; maintenance 1%) with body temperature thermostatically controlled using a heating pad. Rimadyl was used for post-operative analgesia. Under aseptic conditions, the animals received bilateral stereotaxic injections (0.3μl/site) into the SCN (±0.25mm medio-lateral to Bregma, 5.5mm deep to dural surface) of an AAV-1 vector encoding pCry1-Cry1::EGFP (3.26×10^12^cg/ml) for circadian expression of Cry1::EGFP fusion, driven by its minimal promoter (SCN^Cry1^, n =9), or pCry1-EGFP control (SCN^Con^, n =9). At the same time, a telemetric transmitter (TL11M2-F20-EET, Data Sciences International, St Paul, MN) connected to electrodes for continuous electroencephalography (EEG) and electromyography (EMG) recordings was implanted subcutaneously. Two screws were implanted above the dura (+1.5mm anterior to Bregma and +1.7mm lateral to Bregma, the second +1.0mm anterior and +1.7mm lateral to Lambda) around which the electrodes for measuring the EEG were placed and secured using dental cement (RelyX Unicem 2 automix; Henry Schein Animal Health, Dumfries, UK). The two EMG leads were inserted into the trapezius muscle ca. 5mm apart and sutured in place (Hasan et al., 2011; Lang et al., 2011). All mice were allowed 10-14 days of recovery before recording EEG/EMG signals. To confirm AAV targetting of the SCN, at the conclusion of the study, mice were culled and the brains dissected, fixed in 4% paraformaldehyde in phosphate buffer, cryopreserved overnight in 20% sucrose in PBS and then sectioned (40μm) on a freezing sledge microtome (Bright Instruments, UK). Confocal microscopy (Zeiss 780 inverted confocal system) of the native EGFP signal in control SCN^Con^ and SCN^Cry1^ groups was used to identify successful targeting of the SCN. Due to inefficient targeting, two of the nine animals were removed from further analysis in the SCN^Cry1^ group and one from the SCN^Con^ group.

### EEG/EMG recordings and determination of vigilance states and spectral analysis

Transmitters were activated on the day before data collection and EEG/EMG were recorded continuously from the freely moving animals in both LD (2 days) and DD (3 days) using Data Sciences International hardware and Dataquest ART v2.3 Gold software (Data Sciences International, ST Paul, MN). Vigilance states for consecutive 4s epochs were classified by visual inspection according to standard criteria: wakefulness (high and variable EMG signal, low-amplitude EEG signal), NREMS (high EEG amplitude dominated by slow waves, low EMG), and REMS (low EEG amplitude, theta oscillations and muscle atonia). Vigilance states were analysed offline using Neuroscore Software (v.2.1 Data Sciences International) with the EEG and EMG signals modulated with a high-pass (3dB, 0.5Hz) and a low-pass (50Hz) analogue filter and manually assessed. For both LD and DD conditions, continuous recordings were analysed and time spent in each vigilance state was expressed as a percentage of the total recording time over various intervals (1h to 24h). All DD recordings were started after at least 7 days of constant conditions. The mean duration of individual bouts of vigilance states was analysed for the 12h light/subjective day and 12h dark/subjective night periods, and between ZT6-12 on baseline day and following 6h of sleep deprivation (SD). The total amount of NREMS during SD was calculated as well as the latency to the first >10 epochs (40 seconds) of NREMS post 6h of SD. Two mice from the SCN^Con^ group were excluded from the sleep analysis (n=1 LD and DD; and n=1 from DD only) due to the lack of sleep-wake data. Spectral analysis was computed for consecutive 4s epochs by a fast-Fourier transform (frequency range: 0.5 -49.80Hz; resolution 0.24Hz; Hanning window function) on the EEG signal for wakefulness, NREMS and REMS. Genotypic differences were determined in LD and DD over either a complete 24h period or a complete circadian cycle, respectively, and expressed as a percentage of total EEG power within all vigilance states for each mouse. Spectral activity in each bin for each mouse was also normalised to the mean of WT animals. Epochs containing EEG artefacts were discarded from the analysis. The time course of spectral activity was also computed in 2h bins during LD and DD for delta (1 -4Hz) during NREMS and post 6h SD and calculated as a percentage of the mean 24h baseline for each mouse.

### Sleep deprivation and novel object recognition test

Mice were recorded for a 24h baseline day followed by 6h SD and a further 18h recovery sleep. SD (ZT0-6) involved gentle procedures, i.e., introduction of novel objects such as nesting material, “fun tubes” and an initial cage change. Novel object recognition was tested in dim red light (<10lux) between ZT20 and ZT22, in a red Perspex box measuring 50×50×50 cm with an overhead camera (Logitech Carl Zeiss Tessar HD 1080P) placed above the arena. The mice were habituated to the arena without objects for 10 min, followed by an initial familiarisation session 24h later where they were exposed to two identical objects for 10 min (plain or patterned Perspex objects e.g., square, pyramid, oval, egg-cup all of similar sizes). After 24h, the mice were re-tested with one of the objects being replaced by a novel object of similar size. Animals are assumed to have remembered the familiar object if they spend significantly more time investigating the novel object during the test phase. The percentage time the animals spent exploring each object in both the familiarisation and test sessions were analysed offline from the video recordings (using software designed by the laboratory of Prof W. Wisden, Imperial College, London, UK (Yu et al., 2014) with the experimenter blind to the genotype of the animal.

### Experimental Design and Statistical Analysis

Analyses were conducted in Prism version 8.0 for macOS X (GraphPad software). One or two-way ANOVA, with repeated measures where relevant, and with *post-hoc* Tukey’s, Dunnett’s or Sidak’s multiple comparisons were used to compare changes in sleep/wake parameters across genotypes. Data from the running-wheels were analysed using ClockLab (Actimetrics Inc., USA), running within Matlab (Mathworks, USA). Circadian period (chi-squared periodogram analysis) and mean DD activity profiles were calculated for each animal, where activity was averaged over 8-10 days of activity and organized into 0.1h bins. Comparisons of sleep-wake bouts, duration and frequency were made using ANOVA between genotypes and a paired two-tailed Student’s t-tests within genotype. In all cases, the experimental unit was an individual mouse. Male mice were used to avoid the confounding effect of oestrus cycles on circadian behaviour patterns observed in female mice. Following attrition due to technical difficulties, the three treatment-group sizes were WT n =5; SCN^Con^ n =7 and SCN^Cry1^ n =7. Given the variance of our measures, these sample sizes would yield a statistical power of 90-95% (G∗Power 3.1, University of Dusseldorf, Germany).

## Results

### Local expression of Cry1 in the SCN initiates circadian wheel-running behaviour

Local, bilateral AAV-mediated nuclear expression of Cry1::EGFP in the SCN of *Cry1/Cry2*-null mice was evident from the EGFP tag (Figure 1A, B, C). It was limited to the SCN and the immediately surrounding hypothalamus, in some cases extending dorsally into the anterior hypothalamic area and paraventricular nucleus (PVN) and/or posteriorly to the retrochiasmatic area and/or anteriorly to the medial preoptic area. There was, however, no consistent pattern of extra-SCN expression between animals. WT mice exhibited robust circadian cycles of wheel-running behaviour, whereas *Cry1,2*-null mice targetted with EGFP (SCN^Con^) in DD were arrhythmic before and after surgery (Figure 1D). In contrast, previously arrhythmic SCN^Cry1^ mice exhibited robust circadian wheel-running behaviour following expression of Cry1::EGFP in the SCN (Maywood et al., 2018). The mean post-surgery activity profile between groups shows the significant circadian rhythmicity in both the wild-types and SCN^Cry1^ mice, whereas activity in SCN^Con^ mice was distributed evenly across the circadian cycle, phase-referenced to the prior LD cycle (Figure 1E) (2xANOVA: Interaction F_45,630_ =4.0 p<0.0001; Time: F_45,630_ =10.8 p<0.0001). Furthermore, the period of 24.8 ±0.2h (n =7) (Figure 1F) in SCN^Cry1^ mice was significantly longer than in WT (n =5; 24.1 ±0.1h; t_10_ =4.03; p<0.005 2-tailed unpaired Student’s t-test) and diagnostic of a Cry1-driven TTFL (van der Horst et al., 1999) (Maywood et al., 2018). Finally, non-parametric analysis of locomotor activity confirmed the excellent circadian organisation of WT mice, and its disorganisation in SCN^Con^ mice, which had low relative amplitude and stability and high variability (Figure 1G, H, I). In contrast, previously arrhythmic SCN^Cry1^ mice showed robust circadian behaviour after surgery, with significantly improved organisation comparable to WT mice (SCN^Cry1^ paired t-test pre- and post-surgery: Relative amplitude: t_6_ =18.6, P<0.0001; Interdaily stability: t_6_ =3.3, P<0.02), and a non-significant trend for lower intra-daily variability).

**Figure 1:**
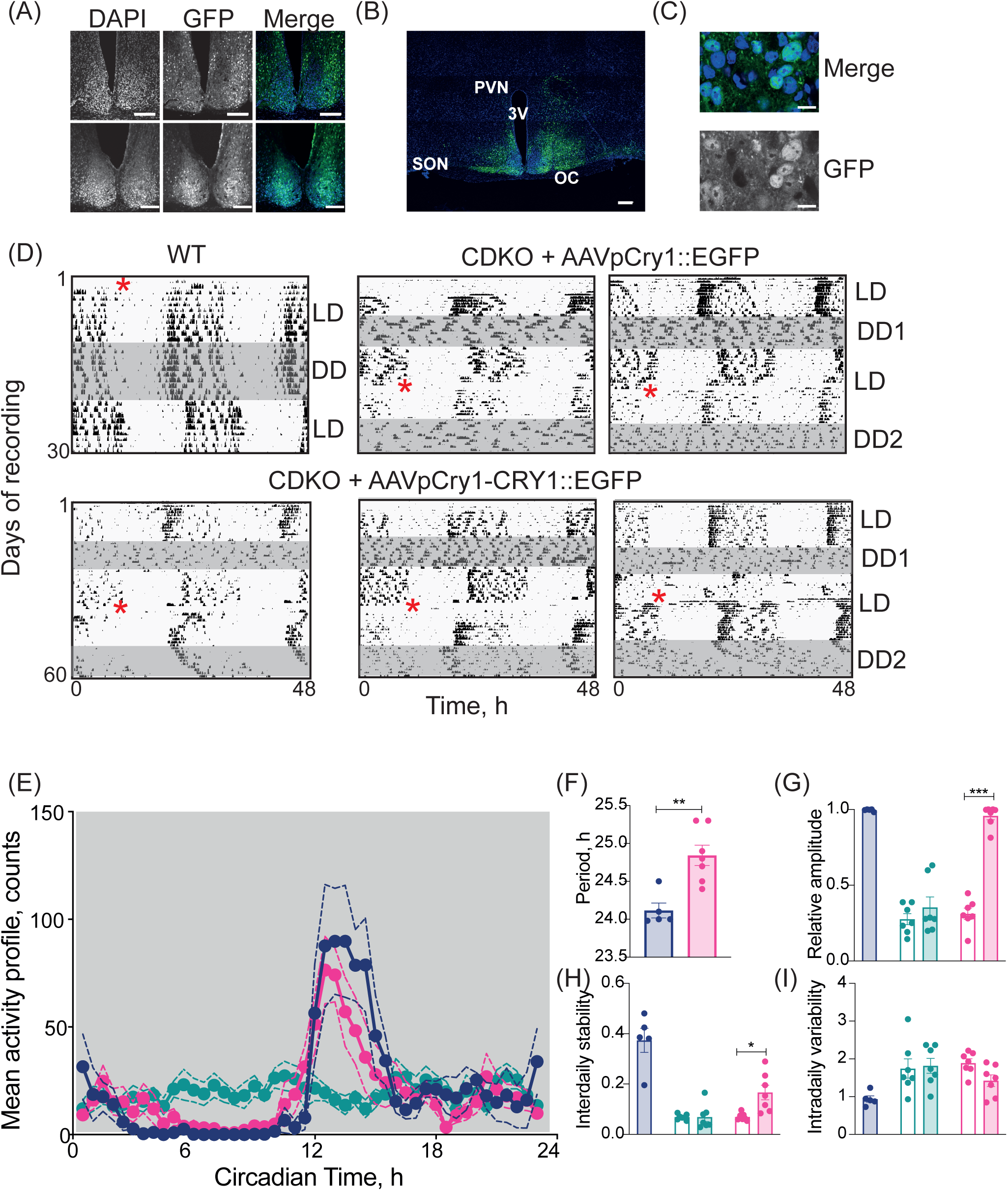
Local expression of Cry1 in the SCN initiates circadian wheel-running behaviour. (A) Fluorescence confocal images from the brains of two mice injected with an AAV pCry1-Cry1::EGFP to restore Cry1 into the SCN (SCN^Cry1^), x20, scale bars =150μm. (B) Tiled (4×4) fluorescence confocal image from the same animal as in the upper panel of (A), x20, scale bar =150μm. (C) High power fluorescence confocal image from the same animal as in the upper panel of (A), showing nuclear co-localisation of EGFP and DAPI signals, x63, scale bar =10μm. (D) Double-plotted wheel-running traces from a wild-type (top left), two SCN^Con^ (top middle and right) and three representative traces from SCN^Cry1^ mice (bottom). Grey shaded areas represent constant darkness (DD) before (DD1) and after (DD2) surgery (denoted by the red Asterix). The CDKO mice were arrhythmic pre-surgery, and the SCN^Con^ mice remained arrhythmic post-surgery, whereas SCN^Cry1^ mice showed significant circadian rhythmicity post-surgery. (E) Mean activity (±SEM) counts over a circadian cycle in wild-type (n =5; blue), SCN^Con^ (n =7; green) and SCN^Cry1^ (n =7; magenta). (2xANOVA time v genotype P<0.001). (F) Circadian period (±SEM) of WT and SCN^Cry1^ mice (∗∗p<0.01 unpaired Student’s t-test). Data Mean (±SEM) and individual points. (G -I) Relative amplitude (G), inter-daily stability (H) and intra-daily variability (I), in WT, SCN^Con^ mice and SCN^Cry1^ mice. Open bars are pre-surgery, closed bars post-surgery (∗p<0.05, ∗∗p<0.01, ∗∗∗p<0.0001 paired Student’s t-test). Data Mean (±SEM) and individual points.

### Local expression of Cry1 in the SCN organises circadian sleep/wake patterning

Initiation of circadian control of wheel-running behaviour by AAV-mediated genetic complementation of the SCN made it possible to explore the degree of control to sleep mediated, uniquely, by a molecularly competent SCN. EEG spectral analysis showed that under DD, in the absence of any masking or other effects of light, the different vigilance states exhibited their characteristic neurophysiological features (Supplementary Figure S2-1A, B, C). Moreover, the genotype had no effect on these parameters, confirming that the absence of Cry proteins does not affect the core molecular and neural machinery that generates the states of wakefulness, REMS and NREMS. The total amount of wake did not vary between groups under LD or DD (Figure 2A), but compared with WT controls, both SCN^Con^ and SCN^Cry1^ mice exhibited more NREMS under LD, as reported (Wisor et al., 2002) (WT 618.4 ±26.4; SCN^Con^694.2 ±20.; SCN^Cry1^ 692.2 ±12.6 min: 1xANOVA F_2,16_=4.7, p=0.024). This likely represents a Cry-dependent trait independent of the SCN and was absent under DD (Figure 2B). On DD, but not LD, SCN^Cry1^ mice showed a small elevation in REMS. Overall, therefore, the genetic manipulations did not have an appreciable effect on the ability of the three groups to exhibit normal vigilance states, especially on DD.

**Figure 2:**
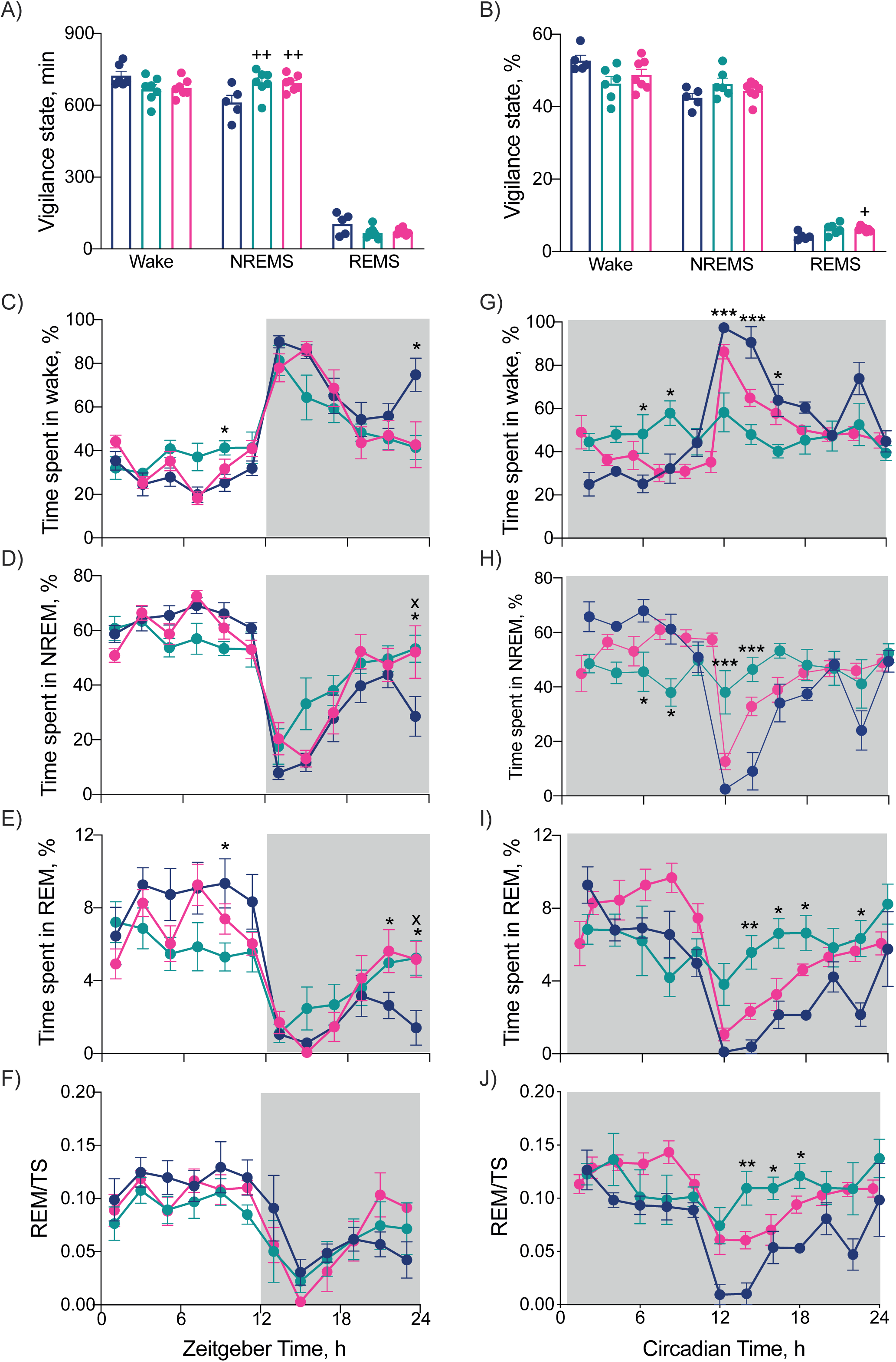
Local expression of Cry1 in the SCN organises circadian sleep/wake patterning. (A, B) Total time spent in wakefulness, NREMS and REMS in (A) 12hL:12D (mins) and (B) in DD (% time) by WT (n =5; blue), SCN^Con^ (n =7; green) and SCN^Cry1^ (n =7; magenta) mice (2xANOVA: P<0.001 post-hoc Tukey’s multiple comparisons test wild-type + p<0.05, ++ p<0.01 vs WT). Mean (±SEM) and individual points. (C-F) Entrained profiles of wakefulness (C), NREMS (D), REMS (E) and REMS/Total sleep (TS; F) in WT, SCN^Con^ and SCN^Cry1^ mice under a 12h light:12h dark photoschedule. Plotted in 2h bins, mean ±SEM. (G-J) Circadian hour profiles of wakefulness (G), NREMS (H), REMS (I) and REMS/Total sleep (TS; J) in WT (n=5; blue), SCN^Con^ mice (n=6; green) and SCN^Cry1^ (n=7; magenta) under DD. Plotted in 2h bins, mean ±SEM. (2xANOVA post-hoc Tukey’s multiple comparison test ∗, ∗∗, ∗∗∗ P<0.05, 0.01, 0.001 WT vs SCN^Con^; x, xx P<0.05, <0.01 WT vs SCN^Cry1^; + P<0.05 SCN^Con^ vs SCN^Cry1^).

We then examined the temporal distribution of sleep/wake. Under LD, WT mice showed appropriate nocturnal wakefulness and more NREMS and REMS in daytime (Supplementary Figure 2-2A, C, E) (WT, LD: 2-tailed Student’s t-test: Wake: t_4_ =24.4, p<0.0001; NREMS: t_4_ =13.9, p<0.0005; REMS: t_4_ =4.2, p=0.013). Equally, both CDKO groups, SCN^Con^ and SCN^Cry1^, had clearly defined day/night differences in wake, NREMS and REMS on LD (2-tailed Student’s t-test: SCN^Con^: Wake: t_6_ =5.0, p<0.005; NREMS: t_6_ =5.4, p=0.0016; REMS: t_6_ =3.2, p=0.018; SCN^Cry1^: Wake: t_6_ =9.7, p<0.0001; NREMS: t_6_ =8.9, p<0.0001; REMS: t_6_ =8.0, p<0.0002) (Supplementary Figure 2-2). Comparison across genotypes nevertheless revealed genotype x time interactions for both wake and NREMS, which exhibited a stronger daily pattern in WT mice (2xANOVA: wake: Interaction F_4,32_ =10.8, p<0.0001; Time: F_2,32_ =904, p<0.0001; Genotype: F_2,16_ =1.8, p =0.2; NREMS: Interaction F_4,32_ =6.2, p<0.0005; Time: F_2,32_ =931, p<0.0001; Genotype: F_2,16_ =4.4, p=0.029).

Under DD, WT mice again showed clear differences between subjective day (higher levels of NREMS and REMS) and night (more wake) (Supplementary Figure 2-2B, D, F) (2-tailed paired Student’s t-test: Wake: t_4_ =17.2, p<0.0001; NREMS: t_4_ =12.4, p<0.0002; REMS: t_4_ =10.9, p<0.0005). In contrast, under DD, the SCN^Con^ mice showed no significant circadian patterning to vigilance states (2-tailed paired Student’s t-test: Wake: t_5_ =0.18, p=0.86; NREMS: t_5_ =0.08, p =0.93; REMS: t_5_ =0.02, p =0.98) (Supplementary Figure 1E). Unlike in LD, where the light imposed a level of organisation on the sleep/wake profiles, the SCN^Con^ mice spent significantly less time in NREMS in the circadian day and significantly more in circadian night, compared to WT. In contrast, SCN^Cry1^ mice showed robust circadian organisation to the sleep-wake cycle, with clear subjective day and night differences comparable to those of WT controls (2-tailed paired Student’s t-test: Wake: t_6_ =7.1, p<0.0005; NREMS: t_6_ =5.4, p<0.002; REMS: t_6_ =6.0, p<0.001). The loss of circadian organisation to sleep in SCN^Con^ mice and its establishment *de novo* in SCN^Cry1^ mice was reflected in significant genotype x time interactions for all three vigilance states (2xANOVA: wake: Interaction F_4,32_ =35.7, p<0.0001; Time: F_1,15_ =117, p<0.0001; Genotype: F_2,15_ =3.2, p =0.07; NREMS: Interaction F_4,30_ =25.3, p<0.0001; Time: F_1,15_ =81, p<0.0001; Genotype: F_2,15_ =2.2, p =0.14; REMS: Interaction F_4,30_ =18.9, p<0.0001; Time: F_1,15_=75.3, p<0.0001; Genotype: F_2,15_ =5.0, p=0.021).

A finer-grained, 2h resolution, analysis of the 24h distribution of sleep/wake emphasised further the effects on sleep-wake patterning of global Cry deficiency and local restoration of clock function in the SCN. Under LD, all groups exhibited a daily pattern of vigilance states, but it had a reduced amplitude in the SCN^Con^ mice, likely reflecting their poor behavioural entrainment to, and/or masking by, the photoschedule (Figure 2C, D, E). Furthermore, in the second half of the light phase, SCN^Con^ mice had significantly more wakefulness (+50min) and less NREMS (−40min) compared with WT mice. Expression of Cry1 in the SCN reversed these deficits (1xANOVA: Wake: F_2,16_ =8.3, p<0.005; post-hoc Tukey’s multiple comparison test: WT v SCN^Con^ p<0.003; WT v SCN^Cry1^ p =0.36; SCN^Cry1^ v SCN^Con^ p<0.035; 1xANOVA: NREMS: F_2,16_ =7.2, p =0.0059; post-hoc Tukey’s multiple comparison test: WT v SCN^Con^ p<0.007; WT v SCN^Cry1^ p =0.55; SCN^Cry1^ v SCN^Con^ p=0.036). Conversely, at the end of the dark phase WT mice showed more wake and less NREMS and REMS that both CDKO groups. Finally, the amount of REM as a proportion of total sleep (TS= NREMS +REMS), which is a measure of circadian control independent of changes in the absolute amount of wakefulness, was high in day and low at night in WT mice and this pattern was replicated by SCN^Con^ and SCN^Cry1^ (Figure 2F). Under LD, therefore, loss of Cry proteins altered the temporal distribution of vigilance states, and this was corrected to some degree by Cry1 expression in the SCN.

The differences between WT and SCN^Con^ mice were amplified in DD. Whereas WT exhibited robust circadian patterning, SCN^Con^ mice failed to show any significant organisation of wake, NREMS or REMS across the circadian cycle (Figure 2G, H, I). Similarly, in WT mice the amount of REM sleep as a proportion of total sleep was highly circadian under DD, whereas it was not in SCN^Con^ mice (DD: WT:1xANOVA F_11,48_ =5.8, p<0.001; SCN^Con^: F_11,60_ =0.9, n.s.) (Figure 2J). These differences were evident in the individual mice in the two groups (Supplementary Figure 2-3A, B). Expression of Cry1 in the SCN had a marked restorative effect on sleep/wake patterns in DD. The SCN^Cry1^ mice showed a WT-like organisation of the sleep/wake cycle across circadian day and night (Figure 2G, H, I), although they did have less wakefulness (ca. 12%) and more NREMS (ca. 10%) and REMS (ca2%) in the circadian night compared with WT. Levels of REMS/TS were higher overall in SCN^Cry1^ mice, but they nevertheless showed a significant circadian rhythm in the distribution of REMS/TS (Figure 2J) and, as with the WT mice, had significantly lower overall levels of REMS/TS in circadian night than did the SCN^Con^ mice (2xANOVA Interaction F_4,32_ =6.6, p<0.001; post-hoc Tukey’s multiple comparison test: WT v SCN^Con^ p<0.005; SCN^Cry1^ v SCN^Con^ p=0.025). This circadian organisation was evident in all individual SCN^Cry1^ mice (Supplementary Figure 2-3C). Together, the LD and DD analyses confirm the interaction between light and the circadian system in organising sleep/wake cycles (Tsai et al., 2009), and suggest that the SCN has a sleep-promoting/wake-suppressing effect in the second half of the LD cycles. Nevertheless, in the absence of a lighting cycle, a molecularly competent SCN is able to impose a circadian distribution of sleep-wake patterns in an otherwise clockless mouse.

### Local expression of Cry1 in the SCN consolidates sleep/wake architecture

Loss of circadian patterning to sleep-wake in SCN^Con^ mice and its restoration in SCN^Cry1^ mice were clear-cut indicators of the autonomous power of the SCN clock. We then examined its effect on sleep/wake architecture, as there was no *a priori* reason to expect that a functional SCN, alone, could reinstate a WT-like structure. Consolidated cycles contain longer and/or fewer bouts for each vigilance state, and fragmented sleep-wake is reflected by shorter and more frequent bouts. When entrained to an LD cycle, WT mice showed a longer duration of wake in the dark phase, and correspondingly fewer episodes of NREMs and REMS at night and more in the light phase (with no systematic changes in their duration) (Figure 3A, B). In comparison, SCN^Con^ mice showed weaker consolidation. They did not exhibit longer wake bouts at night than at day, and nocturnal wake bouts were shorter than in WT mice, with NREMS and REMS bouts longer, as reported previously for CDKO mice (Wisor et al., 2002). Nevertheless, bouts of NREMS and REMS were more frequent in the light phase, as in WT mice. Local expression of Cry1 in the SCN rescued the deficits of SCN^Con^ mice: the duration of wake bouts was significantly longer at night, and no different from WT measures (2-tailed paired t-test: Wake: WT: t_4_ =8.8, p<0.0005; SCN^Con^: t_6_ =1.3, p=0.3; SCN^Cry1^: t_6_ =4.3, p<0.005), the duration of nocturnal NREMS and REMS bouts was comparable to WT mice, and there were fewer bouts of nocturnal NREMS and REMS (LD: 2-tailed paired t-test: NREMS: WT: t_4_ =5.2, p=0.0063; SCN^Con^: t_6_ =4.2, p=0.0054; SCN^Cry1^: t_6_ =8.6, p<0.0001; REMS: WT: t_4_ =5.2, p=0.0066; SCN^Con^: t_6_ =3.6, p=0.011; SCN^Cry1^: t_6_ =7.6, p<0.0005).

**Figure 3:**
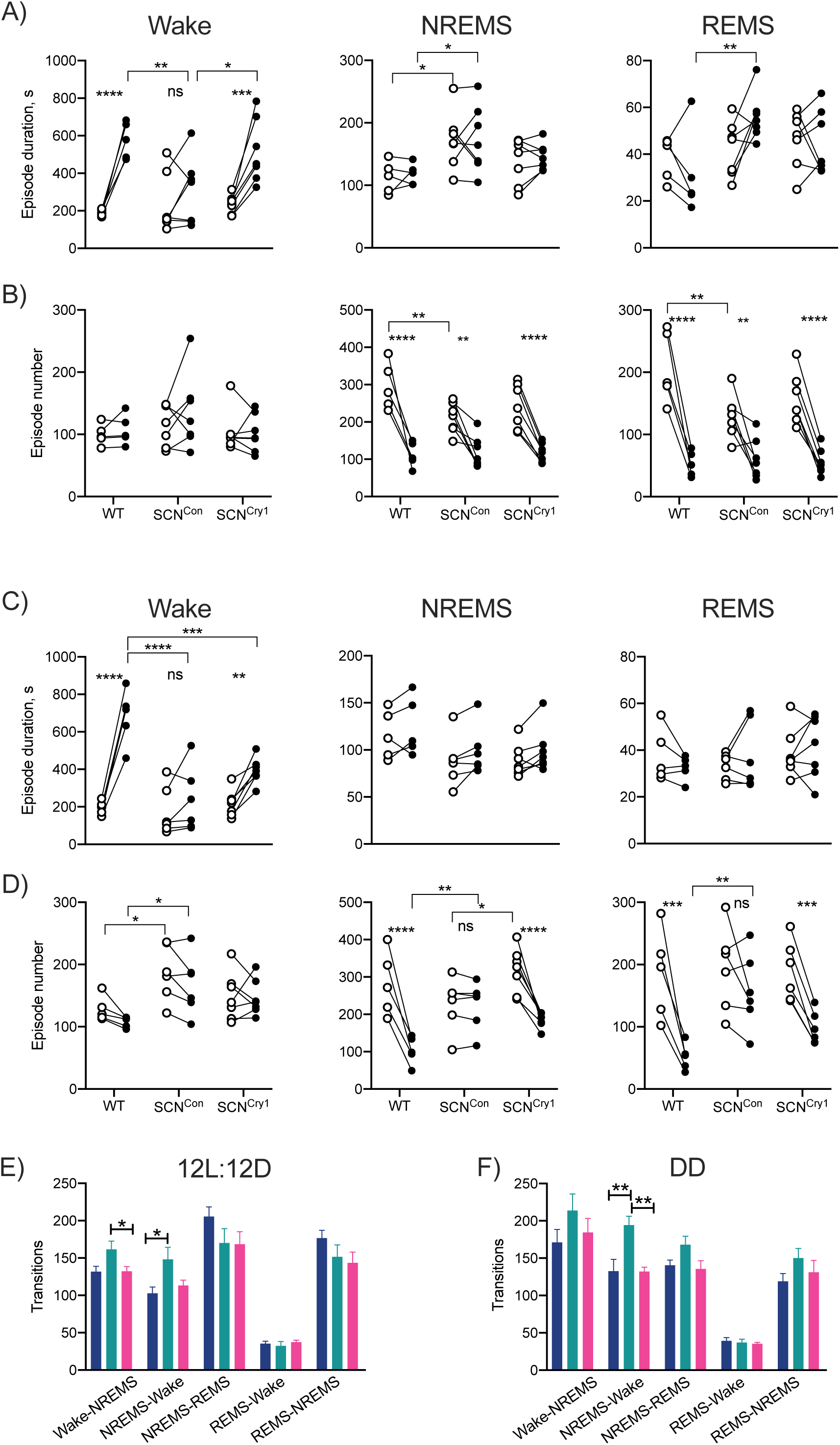
Consolidation of sleep/wake architecture by local expression of Cry1 in the SCN. (A) Duration of bouts in wakefulness, NREMS and REMS in light phase (open circles) and dark phase (closed circles) in WT (n =5), SCN^Con^ (n =7) and SCN^Cry1^ (n=7) mice in entrained (LD) conditions. (B) Mean number of bouts of wakefulness, NREMS and REMS in light phase (open circles) and dark phase (closed circles) of WT, SCN^Con^ and SCN^Cry1^ mice in LD. (C) Duration of bouts of wakefulness, NREMS and REMS in subjective day (open circles) and subjective night (closed circles) in WT, SCN^Con^ and SCN^Cry1^ mice in free-running conditions (DD). (D) Mean number of bouts of wakefulness, NREMS and REMS in subjective day (open circles) and subjective night (closed circles) WT, SCN^Con^ and SCN^Cry1^ mice in DD. (2xANOVA with post-hoc Tukey’s multiple comparison test ∗, ∗∗, ∗∗∗, ∗∗∗∗ P<0.05, <0.01, <0.001, <0.0001) (Paired t-tests show light/subjective day vs dark/subjective night difference within genotype; ∗, ∗∗, ∗∗∗P<0.05, <0.01, <0.001). (E) Mean (±SEM) number of transitions between sleep-wake states in LD. (F) Mean (±SEM) number of transitions between sleep-wake states in DD. (1xANOVA with post hoc Tukey’s multiple comparisons test ∗, ∗∗ P<0.05, 0.01).

The differences between groups in sleep consolidation were even more stark under circadian free-running conditions. WT mice retained their longer duration of nocturnal wake bouts (t_4_ =9.0, p<0.001) and more bouts of NREMS (t_4_ =6.0, p<0.005) and REMS (t_4_ =4.8, p=0.0085) in circadian daytime (Figure 3C, D). SCN^Con^ mice, however, exhibited no significant differences in the duration or number for any vigilance state (Wake duration: SCN^Con^: t_5_ =1.4, p=0.2; NREMS number: SCN^Con^: t_5_ =0.8, p=0.5; REMS number: SCN^Con^: t_5_ =1.4, p=0.2). Bouts of wake in circadian night were significantly shorter than in WT mice, and nocturnal bouts of all three states were more numerous than in WT mice, reflecting the loss of consolidated wake in circadian night. Equally, SCN^Con^ exhibited significantly more episodes of wake in circadian daytime than did WT mice. Importantly, all of these deficiencies in SCN^Con^ mice were corrected by expression of Cry1 solely in the SCN. In SCN^Cry1^ mice, nocturnal wake bouts were significantly longer than in subjective day (t_6_ =4.8, p<0.005), albeit not as long as in WT mice, and bouts of NREMS (t_6_ =6.4, p<0.001) and REMS (t_6_ =4.5, p<0.005) were significantly more numerous in circadian day than in circadian night. Thus, global loss of Cry proteins destabilises sleep-wake structure on LD and even more so under DD, but a molecularly competent SCN is sufficient to correct this deficit and impose WT-like consolidation in clockless mice.

These differences in sleep-wake consolidation between groups were emphasised further by the number of transitions between wake-NREMS and NREMS-wake. In both LD and DD, SCN^Con^ mice showed more transitions than did WT mice, and this was reversed in SCN^Cry1^ mice (Figure 3E, F) (1xANOVA: LD: Wake-NREMS: F_2,16_ =5.8, p=0.012; Tukey’s multiple comparisons test WT v SCN^Con^ p=0.033; SCN^Con^ v SCN^Cry1^ p=0.021: NREMS-wake: F_2,16_ =5.5, p=0.0148; Tukey’s multiple comparisons test WT v SCN^Con^ p=0.0203; SCN^Con^ v SCN^Cry1^ p=0.0481) and DD compared with WT and SCN^Cry1^ mice (1xANOVA: DD: Wake-NREMS: F_2,15_ =1.1, p =0.3; NREMS-wake: F_2,15_ =10.8 p<0.005; Tukey’s multiple comparisons test WT v SCN^Con^ p=0.0044; SCN^Con^ v SCN^Cry1^ p=0.0021). This further confirms that rhythmic expression of Cry1 in the SCN of *Cry1/Cry2*-null mice stabilises the sleep-wake over the 24h LD cycle and over circadian time, consistent with the view that the SCN pacemaker drives the maintenance of wakefulness and the consolidation of sleep, as appropriate, across the LD cycle and across subjective day and night.

### Local expression of Cry1 in the SCN influences sleep homeostasis

To what extent can SCN-mediated consolidation of sleep timing and patterning affect homeostasis? Delta power (1-4Hz) during NREMS is a commonly used index of sleep homeostasis, with higher levels indicating increased sleep need. Under LD, NREMS delta power in WT mice declined spontaneously across the inactive phase and increased during the dark phase, coincident with increased wake (1xANOVA: F_3,12_ =15.8, p<0.0005; post hoc Dunnett’s multiple comparisons test vs ZT10: p=0.009 ZT2, p=0.006 ZT18, p=0.03 ZT20) (Figure 4A). This pattern was clearly under circadian regulation in DD (Figure 4B) (1xANOVA: F_11,40_ =20.1, p<0.0001; post hoc Dunnett’s multiple comparisons test vs CT10: p=0.019 CT2, p<0.0001 CT14-20, p=0.0028 CT22, p=0.0032 CT24). In contrast, SCN^Con^ mice revealed only a low amplitude pattern on LD (1xANOVA: SCN^Con^: F_4,23_ =3.9, p=0.0165), and no significant circadian pattern in DD (Figure 4B) (1xANOVA: F_11,58_ =0.78, p=0.66). SCN^Cry1^ mice had a significant rhythm in delta power not only in LD but also in DD (1xANOVA: LD: F_11,80_ =14.5, p=0.0001, post hoc Dunnett’s multiple comparisons test vs ZT10 p=0.015 ZT16, p =0.058 ZT18, p=0.0048 ZT20; DD: F_11,80_ =4.7, p<0.0001, post hoc Dunnett’s multiple comparisons test vs CT10: p =0.005 CT12, p=0.0005 CT13.94, p=0.0007 CT15.88, p=0.046 CT17.82, p=0.026 CT19.76). In LD, the peak amplitude of EEG delta power of SCN^Cry1^ mice was the same as in WT, and although in DD the peak amplitude was reduced compared to WT, it was nevertheless appropriately phased (Figure 4A, B). This lower peak may reflect the slightly higher levels of NREMS in the early circadian dark phase in DD compared to WT, a difference which was not evident in LD (Figure 3D, H). Finally, all groups showed a similar latency to first NREMS episode >100s on the dark-to-light transition (WT =18.9 ±8.9min, n =5; SCN^Con^ =18.0 ±6.8min, n =7; SCN^Cry1^ =17.5 ±5.2min, n =7; 1xANOVA: F_2,16_ =0.01, p=0.99). Together these data demonstrate that rescuing Cry1 in the SCN restores rhythmic expression of NREMS delta power in both LD and DD, confirming the appropriate phasing and organisation of sleep-wake across the 24h/circadian cycle. SCN-mediated circadian organisation and consolidation were therefore accompanied by appropriate dynamic signalling of sleep need.

**Figure 4:**
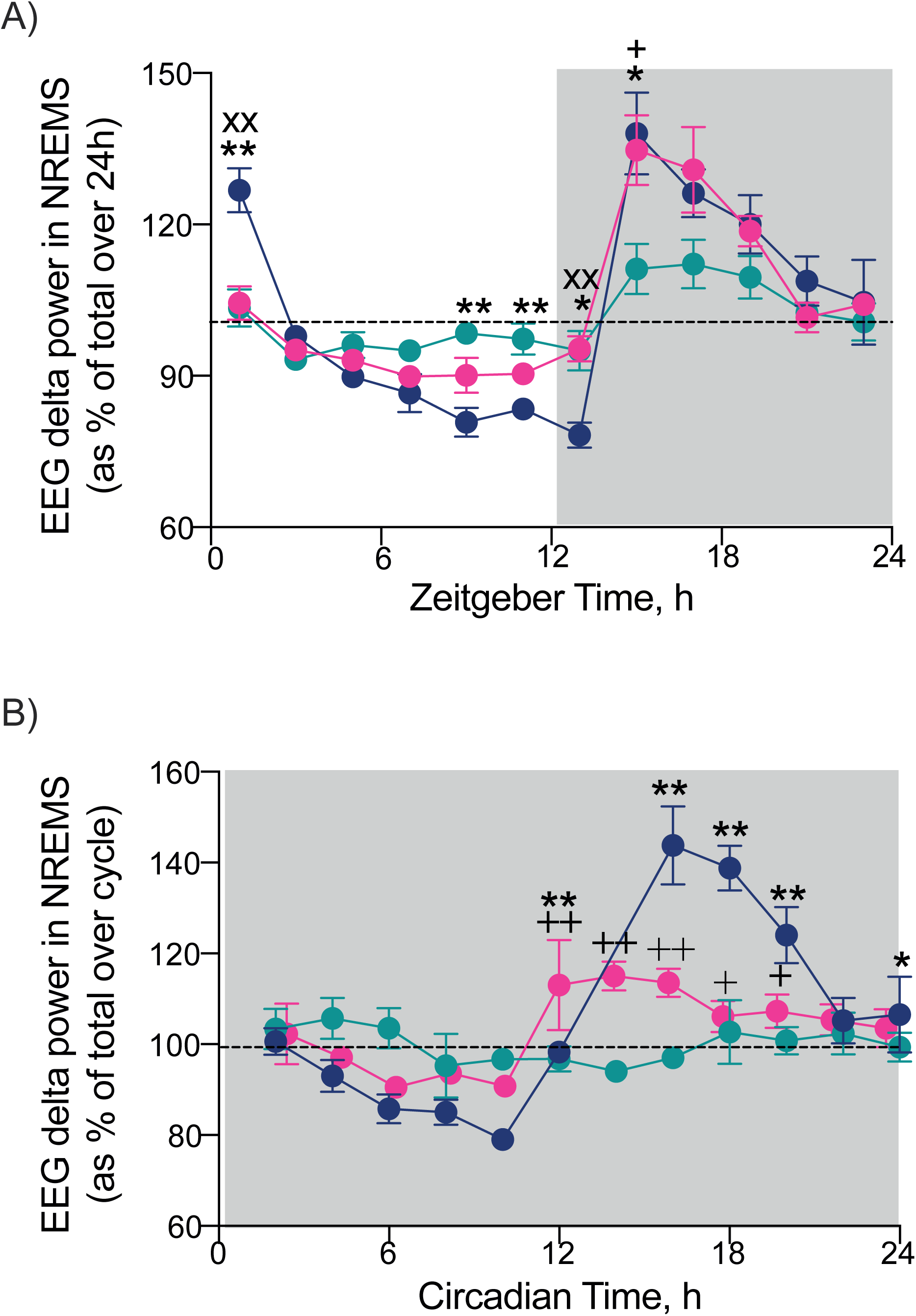
Local Cry1 expression in the SCN affects NREMS delta power (1-4Hz) under entrained and free-running conditions. (A) NREMS EEG delta power in WT (n=5; blue), SCN^Con^ (n=7, green) and SCN^Cry1^ (n=7, magenta) mice in entrained by LD. Plotted in 2h bins, mean ±SEM. (2xANOVA post-hoc Sidak’s multiple comparison test ∗, ∗∗, P<0.05, <0.01, WT vs SCN^Con^; + P<0.05 SCN^Con^ vs SCN^Cry1^; xx P<0.01 WT vs SCN^Cry1^). (B) NREMS EEG delta power in WT, SCN^Con^ and SCN^Cry1^ mice free-running in DD. Plotted in 2h bins, mean ±SEM). (1xANOVA shows WT and SCN^Cry1^ mice have significant rhythms ∗, ∗∗ P<0.05, 0.01 WT vs circadian time (CT) 10; +, ++ P<0.05, 0.01 SCN^Cry1^vs CT10.06. There was no significant rhythm in control treated mice).

Having shown that the SCN alone can direct the circadian patterning and stabilisation of sleep-wake cycles, we next tested whether there was any SCN-dependent interaction between the circadian and homeostatic processes by measuring the EEG responses to 6h sleep deprivation (SD) starting at lights on. SD was equally effective across the three groups, with no significant differences in the small amount of NREMS (WT 9.8 ±3.5min, SCN^Con^ 13.6 ±2.7min, SCN^Cry1^ 7.4 ±2.5min, n =5, 7, 7 respectively) or the latency to sleep post-SD (time to first NREMS bout >100s duration, WT 12.9 ±6.8min,; SCN^Con^ 10.4 ±3.4min; SCN^Cry1^ 10.3 ±3.7min). In the 2h immediately after SD, all groups showed a significant and equal increase in delta power when NREMS occurred in that interval (Figure 5A) reflecting greater homeostatic sleep pressure, and with no difference between groups (2xANOVA: Interaction F_2,16_ =0.2, p=0.8; Group: F_2,16_ =1.2, p=0.3; SD: F_1,16_ =83.9, p<0.0001). Genotype did not, therefore, affect the neurophysiological capacity to sense and respond to sleep deprivation during subsequent NREMS when it did occur (Wisor et al., 2002). Furthermore, recovery from SD, compared to baseline, was not different between genotypes, insofar as accumulated sleep loss increased during SD, but then decreased at the same rate in all three groups over the subsequent 18h (Figure 5B). By whatever mechanism, all three groups recovered lost sleep i.e., exhibited homeostasis.

**Figure 5:**
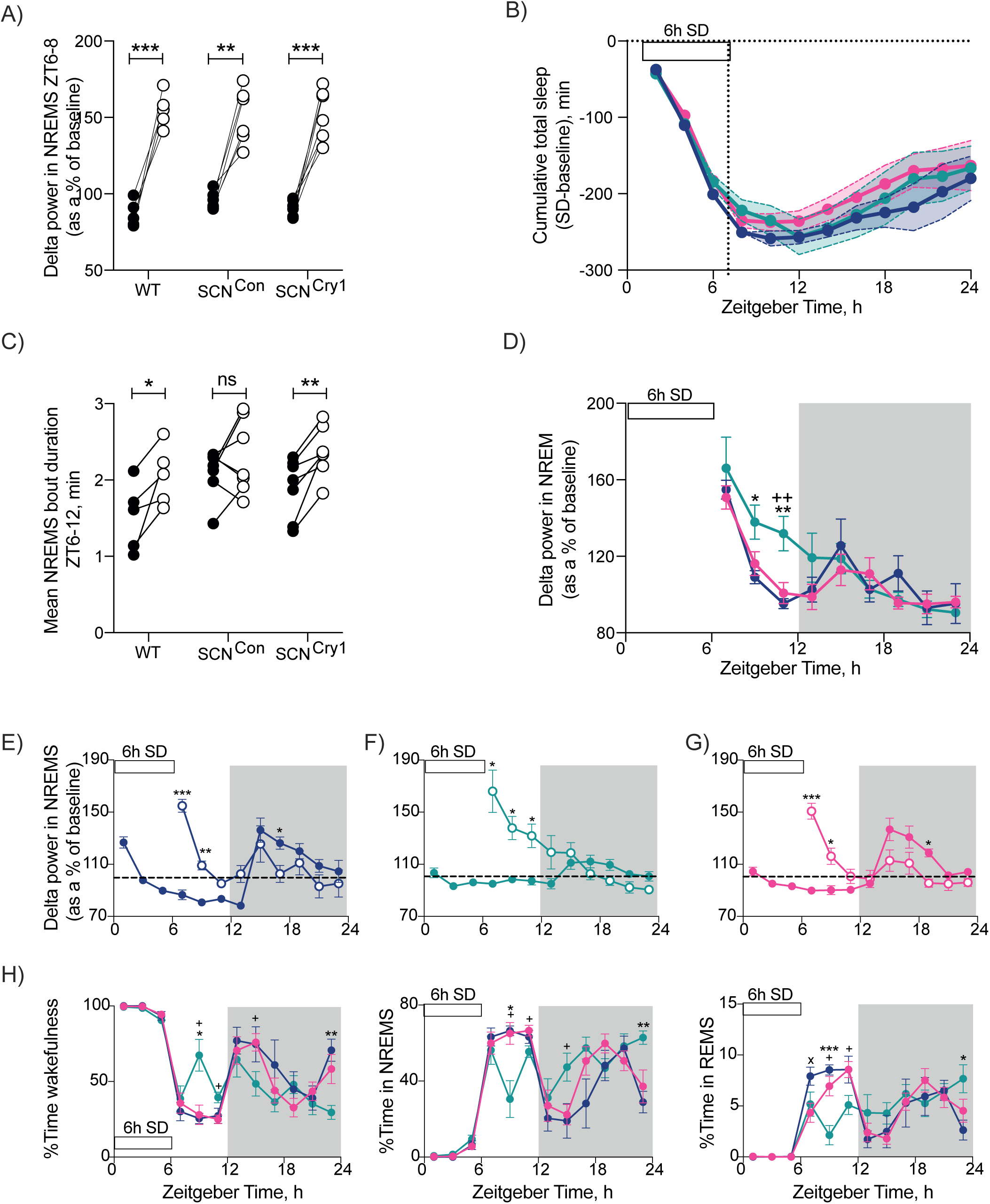
Effect of rescue of rhythmic Cry1 expression in the SCN on the homeostatic response to 6h sleep deprivation. (A) Changes in NREMS delta power (1-4Hz) in the first 2h of recovery sleep (ZT6-8; open circles) compared with baseline sleep (ZT6-8; closed circles) in individual WT, SCN^Con^ and SCN^Cry1^mice (Paired t-tests show baseline vs 6h SD difference within genotype; ∗∗, ∗∗∗P<0.01, <0.001). (B) Cumulative total sleep loss (NREMS + REMS, mean ±S.E.M) over the 6h of SD with subsequent partial recovery during the 18h post-SD in WT (n=5, blue), SCN^Con^ (n=7, green) and SCN^Cry1^ (n=7, magenta) mice. (ANOVA detected no significant differences between the three groups). (C) Changes in NREMS bout duration between ZT6-12 on baseline day (closed circles) and after 6h SD (open circles) in individual WT, SCN^Con^ and SCN^Cry1^ mice. (∗, ∗∗P<0.05, <0.01 baseline vs 6h SD within genotype, paired t-tests;). (D) Time-course of decline in EEG delta power in NREMS following 6h of SD in WT, SCN^Con^ and SCN^Cry1^mice (mean ±SEM, 2xANOVA post-hoc Tukey’s multiple comparison test ∗, ∗∗ P<0.05, 0.01 WT vs SCN^Con^; ++ P<0.01 SCN^Con^ vs SCN^Cry1^). (E-G) Time-course of EEG delta power during NREMS on baseline day (closed circles) and following 6h SD (open circles; data replotted from panel D) (mean ±SEM, 2xANOVA with post-hoc Tukey’s multiple comparison test ∗, ∗∗, ∗∗∗ P<0.05, 0.01, 0001). (H-J) Percentage of time (mean ±SEM) spent in wakefulness, NREMS and REMS during 6h of sleep deprivation (SD) and 18h of recovery in WT, SCN^Con^ and SCN^Cry1^ mice (2xANOVA post-hoc Tukey’s multiple comparison test ∗, ∗∗, ∗∗∗ P<0.05, 0.01, 0.001 WT vs SCN^Con^; x P<0.05 WT vs SCN^Cry1^; + P<0.05 SCN^Con^ vs SCN^Cry1^).

Notwithstanding overall comparability between genotypes, there were also informative differences. Over the 6h immediately following SD (ZT6-12), WT but not SCN^Con^ mice showed significant increases in the duration of NREMS bouts compared with the equivalent period on the baseline day, a marker of increased sleep need (Figure 5C). The response of SCN^Cry1^ mice was the same as in WT (2-tailed paired t-test WT: t_4_ =3.0, p=0.040; SCN^Con^: t_6_ =1.2, p=0.3; SCN^Cry1^: t_6_ =4.6, p<0.005). The time-course for the decline in delta power after SD was also different. It declined progressively across the light phase in WT mice, but more slowly in SCN^Con^ mice and, unlike in WT mice, did not reach baseline levels during the light phase (Figure 5D, E, F). Expression of Cry1 in the SCN corrected these deficits (Figure 5D, G). Thus, although there were no significant differences in the overall recovery of sleep loss after SD, there were significant differences in its time-course during the light phase (Figure 5H-J). Whereas WT mice showed a sustained absence of wake (Figure 5H) and elevation of both NREMS and REMS in the light phase (Figure 5I, J), the SCN^Con^ mice exhibited significantly less NREMS and more wakefulness. Again, this was rescued in SCN^Cry1^ mice (2xANOVA: NREMS: Interaction F_2,16_ =3.0, p=0.076; Group F_2,16_ =7.74, p=0.0045; post-hoc Sidak’s multiple comparison test: WT p=0.97; SCN^Con^ p=0.0365; SCN^Cry1^ p=0.91; Wake: Interaction F_2,16_ =2.77, p=0.093; Group F _2,16_ =9.33, p=0.0021; post-hoc Sidak’s multiple comparison test: WT p=0.95; SCN^Con^ p=0.029; SCN^Cry1^ p=0.99). This suggests that SCN^Con^ mice had a decreased sleep pressure and/or an inability to maintain consolidated NREMS at this phase of the LD cycle. Given that analysis of NREMS delta power indicated that decreased sleep pressure was not the case in SCN^Con^ mice (Figure 5D, F), it is likely that consolidation may have been limiting, as shown by the failure to express longer NREMS bouts (Figure 5C). Poor consolidation at this phase altered the time course to recovering sleep loss, and may reflect an interaction between the circadian and homeostatic processes regulating sleep-wake at the end of the light phase. Loss of Cry proteins did not, therefore, globally affect neurophysiological mechanisms of homeostatic sleep recovery, but in SCN^Con^ mice with an ineffective SCN clock, the dynamics of recovery were altered and the expression of Cry1 in the SCN corrected this (Figure 5H, I, J) (2xANOVA: Wake: Interaction F_22,176_ =3.05, p<0.0001; Time F_6,89_ =41, p<0.0001; Genotype F_2,16_ =0.76, p=0.49; NREMS: Interaction F_22,176_ =3.01, p<0.0001; Time F_6,89_ =42.3, p<0.0001; Genotype F_2,16_ =1.03, p=0.38; REMS: Interaction F_22,176_ =2.97, p<0.0001; Time F_6,93_ =24.01, p<0.0001; Genotype F_2,16_ =0.11, p=0.9). Cry proteins and a competent SCN clock are not, therefore, necessary components of this fundamental sleep homeostatic mechanism, but they do regulate its time-course.

### Cry1 expression in the SCN alone rescues performance in the novel object test

To determine whether SCN-mediated circadian control over phasing and consolidation of sleep/wake cycles has consequences for brain function, we assessed cognitive performance in the Novel Object Recognition (NOR) task, a recognised sleep-dependent behaviour (Palchykova et al., 2006). There were no statistically significant differences between groups in time spent exploring the objects during training (Figure 6A, B) (2xANOVA Interaction: F_2,15_ =0.2, p=0.8; Object: F_2,15_ =0.2, p=0.7; Group: F_2,15_ =0.8, p=0.4). WT mice demonstrated robust memory for the familiar object by spending significantly more time exploring the novel object when tested (2-tailed paired t-test: t_4_ =16.9, p<0.0001). In contrast, SCN^Con^ mice failed to discriminate between the novel and familiar objects, spending less time exploring the novel object than did WT mice (Figure 6C) (2-tailed paired t-test: t_6_ =1.4, p=0.2). Thus, the global absence of Cry proteins compromised performance in a test of memory known to be sleep-dependent. The initiation of circadian competence in the SCN of SCN^Cry1^ mice, and thereby organisation of sleep/wake cycles, resulted in all 7 SCN^Cry1^ mice showing a highly significant preference for the novel object (2-tailed paired t-test: t_6_ =4.8, p<0.005). These results demonstrate that global loss of Cry1 proteins compromises NOR performance, and that a molecularly competent circadian pacemaker within the SCN alone can establish the necessary organisation and consolidation of sleep/wake, and can also sustain sleep-dependent memory.

**Figure 6:**
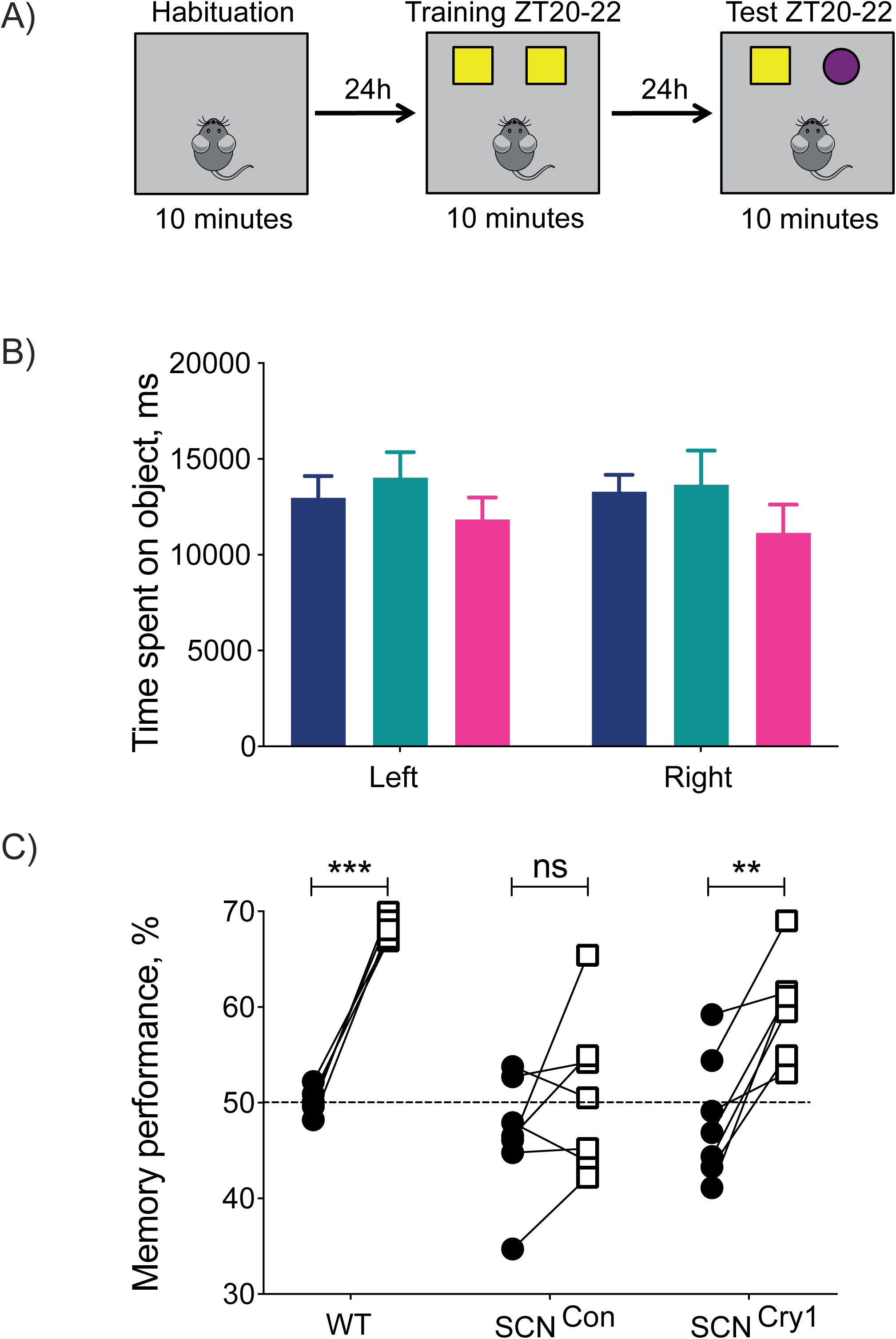
Cry1 expression in the SCN of clockless mice rescues performance in the novel object test. (A) Protocol for the sleep-dependent memory test where, during their nocturnal active phase between ZT20 and ZT22, mice are habituated to the test arena, and 24h later returned and allowed to investigate 2 identical objects. Following a further 24h, one object is replaced with a novel object and the mice are again returned. (B) The amount of time (mean ±SEM) the WT (n= 5, blue), SCN^Con^ (n =7, green) and SCN^Cry1^ (n =7, magenta) mice spent exploring the two objects during the training phase. There were no significant differences between objects and groups (2xANOVA), all mice showing ca. 50% interest. (C) Relative interest of individual mice shown to the two objects during training (close circles) and during testing with a novel object (open circles). Both the wild-type and SCN^Cry1^ groups showed significant preference for the novel object during testing, whereas the SCN^Con^ mice did not (2xANOVA with post-hoc Sidak’s multiple comparisons test ∗∗, ∗∗∗P<0.005, 0.0001 old vs new).

## Discussion

Our approach to address the specific role of the SCN in sleep regulation used genetically clockless *Cry1/Cry2*-null mice that in the absence of LD have no circadian patterning to sleep-wake cycles, poorly consolidated phases of sleep and wake, compromised dynamics of homeostatic recovery sleep, and impaired sleep-dependent memory (van der Horst et al., 1999; Wisor et al., 2002; Wisor et al., 2008; Maywood et al., 2018). We initiated *de novo* circadian rhythmicity locally to the SCN by virally mediated expression of Cry1 (Edwards et al., 2016). The rest of the brain and periphery remained Cry-deficient and circadian-incompetent. As expected, we established circadian locomotor activity rhythms in such SCN^Cry1^ mice (Maywood et al., 2018) comparable to the effect of WT SCN grafts in *Cry1/Cry2*-null mice (Sujino et al., 2003). This allowed us to test the contribution of the SCN molecular clockwork to the temporal regulation of sleep. Genetic rescue of the SCN established sleep-wake cycles that were appropriately phased and consolidated, and accompanied by improved performance in a test of sleep-dependent memory. Furthermore, the data suggest that the circadian system promotes wake in subjective night and facilitates sleep, by promoting its intensity and consolidation, in subjective daytime. Although the neural circuits and neurophysiological processes that sense and direct sleep homeostasis are not compromised by global Cry-deficiency, our data do illustrate an interaction between the homeostatic and circadian mechanisms during the recovery from SD. We conclude that the SCN has continuous influence on sleep-wake organisation across the circadian cycle and, directly or indirectly, modulates sleep consolidation and homeostatic regulation.

Overall sleep-wake distributions across the entrained and free-running cycles showed modest differences between groups, with *Cry1/Cry2*-null mice (SCN^Con^ and SCN^Cry1^) showing an overall increase in the amount of NREMS in LD, as reported (Wisor et al., 2002). Studies using other global circadian mutations/deletions or ablation of the SCN in otherwise intact animals (Naylor et al., 2000; Easton et al., 2004; Laposky et al., 2005; Mistlberger, 2005) have demonstrated either no differences or an increase/decrease in NREMS, making it difficult to interpret whether any phenotypes are due to an extra-SCN circadian effect, or a more global effect on the dynamics of the complex neural circuitry underlining the control of sleep-wake states (Saper et al., 2010). Nevertheless, the SCN^Con^, but not the SCN^Cry1^ mice, did show a significant decrease in the amount of NREMS in the second half of the light/rest phase, suggesting that restoring rhythmicity to the SCN enables promotion of sleep/inhibition of wake at a time when the homeostatic pressure to sleep has declined. Furthermore, whereas the rescued animals showed a WT-like consolidation of wake episodes in the dark/active phase in both LD or DD, the control treated mice did not, and this was reflected in their reduced amplitude or absence of rhythmic time-course for NREMS delta power in LD and DD, respectively. Together these results show the mouse SCN clock has opposing influences on sleep-wake organisation across the cycle: promoting wakefulness in the dark/subjective day and sleep in light/subjective night.

The dynamics of sleep homeostasis were examined following sleep deprivation, to assess whether there is an interaction between the circadian and homeostatic regulatory processes. Evidence suggests influences of sleep homeostasis on the functioning of the circadian clock (Deboer et al., 2003; Deboer et al., 2007; Schmidt et al., 2009), and that these processes can act independently (Tobler et al., 1983; Shiromani et al., 2004). Conversely, global circadian clock mutants can also show altered changes in NREMS EEG delta power in response to SD, consistent with a role for clock genes in sleep homeostasis (Naylor et al., 2000; Wisor et al., 2002; Laposky et al., 2005; Dijk and Archer, 2010; Curie et al., 2013). As discussed above, however, this does not necessarily indicate a role for the SCN in regulating homeostasis. Indeed, *Cry1/Cry2*-null mice, and mice lacking the Cry2 gene (SCN^Cry1^) have an intact sleep homeostatic response (Wisor et al., 2002; Wisor et al., 2008), suggesting that the SCN is not necessary for the expression of the initial neurophysiological response to SD. Nevertheless, in our study SCN^Con^ mice had an altered time-course in their recovery from SD, most notably between ZT6-12 when the SCN is exerting a sleep-promoting influence. This suggests a role, direct or indirect, for the circadian timing system in modulating the recovery from SD, and implies an interaction between the homeostatic and circadian processes, to ensure prolonged and consolidated sleep at a phase when sleep pressure is low under baseline conditions. Studies in humans have similarly postulated a role for the circadian system in influencing sleep homeostatic mechanisms (Lazar et al., 2015), although it remains to be established whether the central circadian clock in the SCN, and/or clocks in other brain areas and/or in the periphery underlie these mechanisms.

How might the SCN exert its effects? The sleep-wake regulatory circuit has been described as a “flip-flop” switch, whereby sleep-wake transitions are regulated by a dynamic interplay of reciprocal inhibition between sleep-promoting and wake-promoting nodes of the hypothalamus and brainstem (Saper et al., 2010). Multiple direct and indirect pathways from the SCN to both sleep- or wake-promoting nodes could therefore influence state-switching over the circadian cycle, which would be expected if the circadian clock has an ongoing active role in regulating both sleep and wake. Activation of lateral hypothalamic (LH) GABA neurons can exert direct synaptic control over the sleep-promoting galaninergic neurons in the ventrolateral preoptic nucleus (VLPO) to promote arousal during NREMS in the light phase (Venner et al., 2019). In addition, a LH-thalamic reticular nucleus (TRN)-GABAergic-thalamocortical inhibitory circuit may be involved in the rapid arousal during NREMS-wake transitions (Herrera et al., 2016). Similarly, the loss of the widely projecting orexin/hypocretin neurons results in more frequent transitions into sleep and so would prevent prolongation of wake episodes (Hara et al., 2001). It may be that changes in GABAergic/glutamatergic drive can affect the rapid changes in state, whereas neuropeptides such as galanin and orexin act as neuromodulators influencing the stabilisation of sleep-wake states, and low (no?) amplitude and/or phasing of output from these cells underlie the sleep phenotypes in *Cry1/Cry2*-null, which are ameliorated following rescue of rhythmicity in the SCN^Cry1^ mice (Willie et al., 2003; Herrera et al., 2016; Venner et al., 2019). In addition, the SCN may also act indirectly because initiation of behavioural rhythmicity in SCN^Cry1^ mice will in turn regulate their metabolic demands, providing feedback from the periphery and/or brain regions. These could in turn influence the timing and homeostatic regulation of sleep (Ehlen et al., 2017; Northeast et al., 2020) by indirect means. For example, overexpression of BMAL1 in skeletal muscle (but not the brain) is reported to influence the daily amount of NREMS, although it did not restore the 24h pattern to sleep/wake, nor the homeostatic responses to SD (Ehlen et al., 2017).

Sleep-dependent memory was severely compromised in SCN^Con^ mice, consistent with other reports of cognitive impairment in *Cry1/Cry2*-nulls (Van der Zee et al., 2008; De Bundel et al., 2013). This could be a result of arrhythmia (in the SCN and/or hippocampal formation), or a non-circadian, molecular consequence of local Cry deficiency. Restoration in SCN^Cry1^ mice refuted the latter, and emphasised the central importance of circadian organisation to cognitive function, be it in the SCN and/or locally in the hippocampal formation, and driven by the SCN. The role of the SCN may, however, be bivalent. In hamsters made arrhythmic using a light pulse paradigm, memory was impaired (Ruby et al., 2008), but the effect was reversed by SCN ablation (Fernandez et al., 2014), suggesting that a dysfunctional SCN signal is more cognitively debilitating than no signal at all. Similarly, in a mouse model of down syndrome, impaired object recognition is restored by SCN ablation (Chuluun et al., 2020). These observations raise the possibility that cognitive deficits might be mitigated by improving circadian amplitude when it is disrupted as, for example, in patients with Alzheimer’s disease (Hatfield et al., 2004) (Leng et al., 2019).

In conclusion, by adopting a gain-of-function approach, we have shown that the SCN clock is not only necessary, but is also sufficient for the temporal control of sleep-wake, facilitating circadian initiation and maintenance of wake, promoting sleep consolidation, the dynamics of homeostatic recovery (but not the ability to sense sleep need), and sleep-dependent memory. Moreover, we show that expression of Cry proteins outside the SCN is not necessary to sustain these processes. Our results therefore add to understanding of the relative contributions of the SCN, extra-SCN clocks and circadian clock genes in the temporal organisation of sleep and wake.

## Acknowledgements

The authors would like to acknowledge the invaluable support of the MRC Laboratory of Molecular Biology Electronics and Mechanical Workshops, particularly Chris Palmer, Tom Pratt in the IT Department and Ares (Biological Services Group). The authors would also like to thank Dr. X. Yu and Prof W. Wisden, Imperial College, London, UK for the use of software for analysing the NOT. This work is supported by MRC core funding to MHH (MC_U105170643).

## Notes

### Competing Interest Statement

The authors have declared no competing interest.

