## Supplementary Figure for "The molecular clockwork of the suprachiasmatic nucleus is sufficient to co-ordinate phasing and stabilisation of sleep-wake cycles and enhance memory deficits in a clockless mouse"

Supplementary Figure 2-1: *Local expression of Cry1 in the SCN organises circadian sleep/wake patterning*

(A, B, C) Relative EEG spectral power (as a percentage of total power) in NREMS (A), wake (B) and REMS (C) in WT (blue), $SCN^{Con}$ (green) and $SCN^{Cry1}$ (red) mice summed across the circadian cycle. Inserts represent the spectral power in percent mean (±SEM) relative to WT (2xANOVA detected no significant differences between groups).

Supplementary Figure 2-2: *Local expression of Cry1 in the SCN organises circadian sleep/wake patterning*

(A, C, E) Total time (mins) spent by WT (blue), $SCN^{Con}$ (green) and $SCN^{Cry1}$ (red) mice in wakefulness (A), NREMS (B) and REMS (C) during the light phase (open) and dark phase (shaded) of entraining LD cycle (mean ±SEM and individual points).

(D, E, F) Time spent (%) by WT (blue), $SCN^{Con}$ (green) and $SCN^{Cry1}$ (red) mice in wakefulness (D), NREMS (E) and REMS (F) during the subjective day (open) and night (shaded) of DD cycle (mean ±SEM and individual points).

(2xANOVA: P<0.001 post-hoc Tukey's multiple comparisons test wild-type +, ++, +++ P<0.05, <0.01, <0.001 vs WT; x, xxx P<0.05, <0.001vs $SCN^{Cry1}$).

(Paired t-tests show light/subjective day vs dark/subjective night difference within genotype; *, **, ***P<0.05, <0.01, <0.001).

Supplementary Figure 2-3: *Local expression of Cry1 in the SCN organises circadian sleep/wake patterning*

Circadian profiles of the time spent (%) in NREMS by individual (A) WT (blue), (B) $SCN^{Con}$ (green) and (C) $SCN^{Cry1}$ (red) mice across the DD cycle. Note the lack of organisation within and between $SCN^{Con}$ mice (B), which was ameliorated in all $SCN^{Cry1}$ mice (C), whose behaviour was similar to individual WT mice (A).

Supplementary FIgure S2-1 (linked with Figure 2)

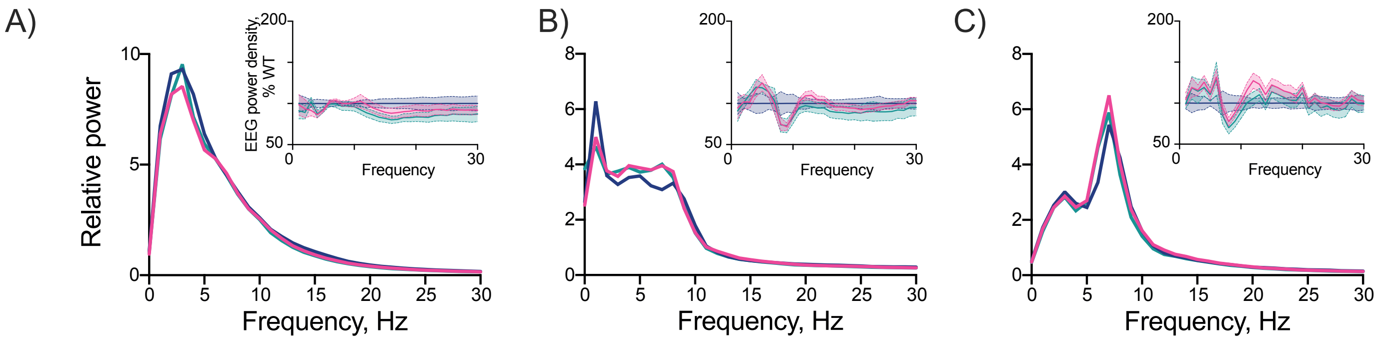

Supplementary Figure 2-2

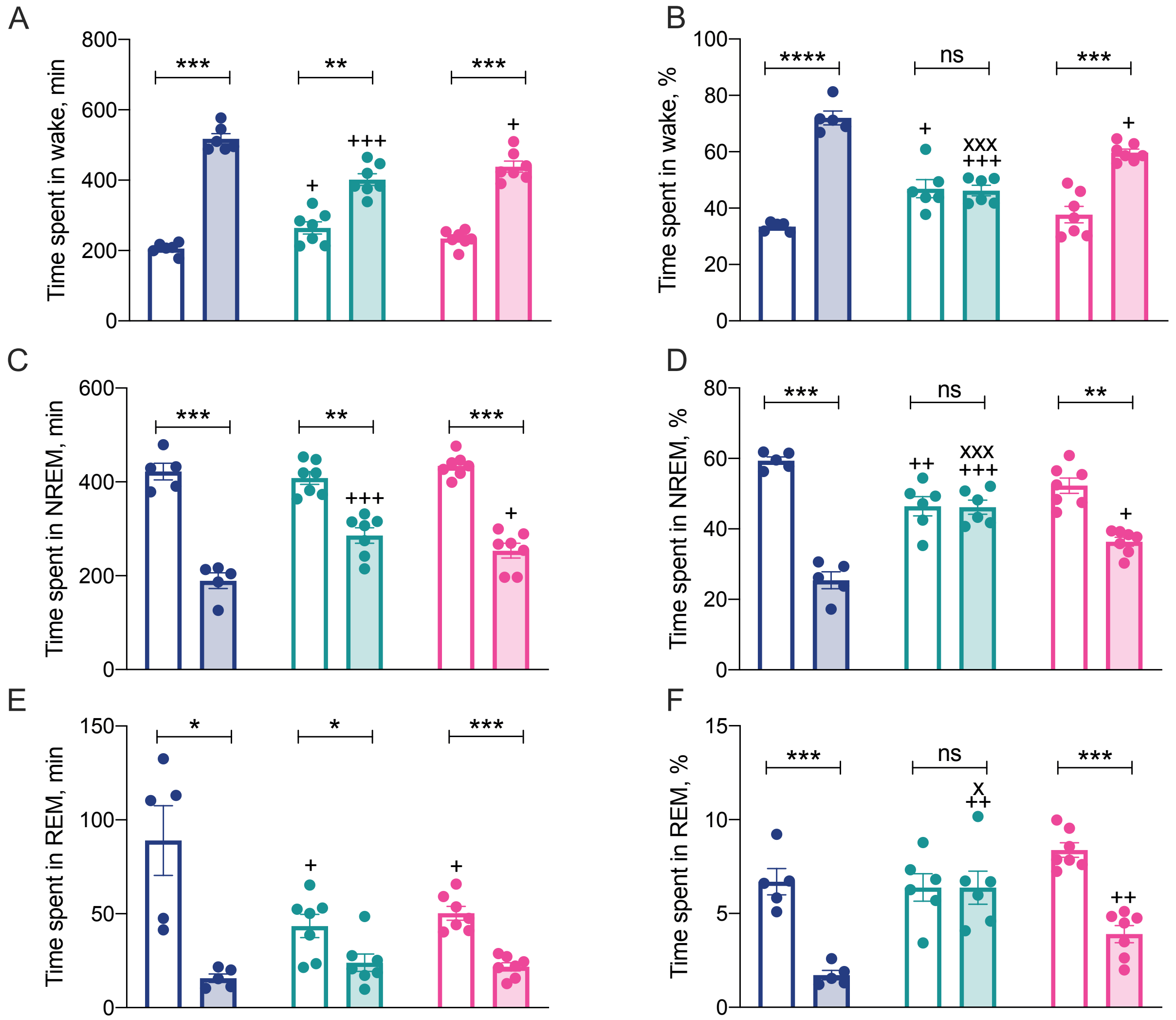

Supplementary Figure 2-3

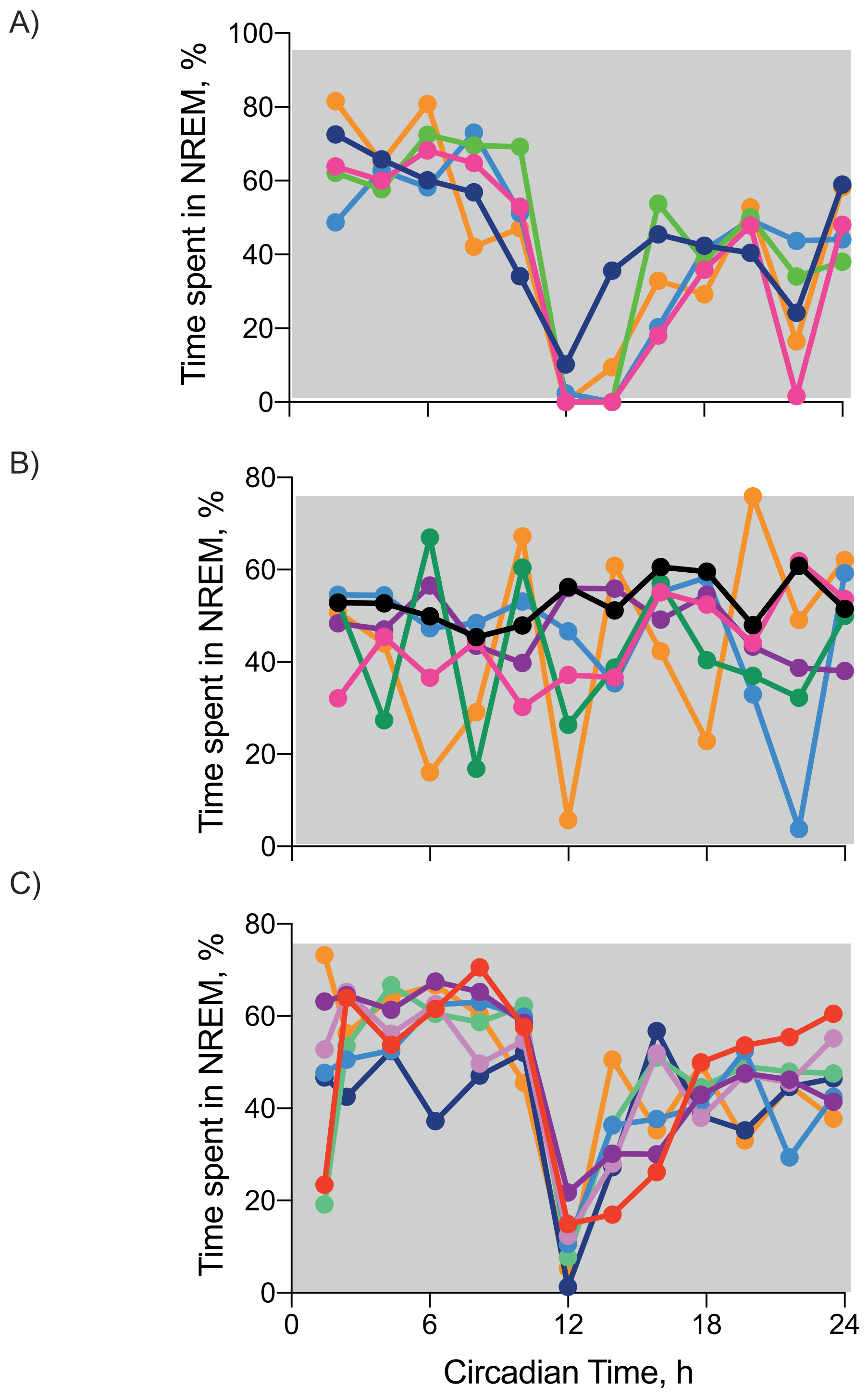
